# On Some Problems of Estimating Fundamental Niche from Physiological Data

**DOI:** 10.1101/716688

**Authors:** Gutiérrez-Ruelas Jesús Salvador, Laura Jiménez, Ana Patricia Quiroz-Reyes, Sabrina Cherizar Sotelo-Pedroza, Soberon Jorge

**Affiliations:** División de Ingeniería, Universidad de Sonora, Blvd. Luis Encinas y Rosales S/N,Col. Centro Hermosillo, Sonora, México; División de Ciencias Exactas y Naturales, Universidad de Sonora, Blvd. Luis Encinas y Rosales S/N,Col. Centro Hermosillo, Sonora, México; División de Ciencias Biológicas y de la Salud, Universidad de Sonora, Blvd. Luis Encinas y Rosales S/N,Col. Centro Hermosillo, Sonora, México; Department of Ecology and Evolutionary Biology and Biodiversity Institute, University of Kansas, 1235 Jayhawk Blvd., Lawrence, KS, USA

## Abstract

Hutchinson proposed the idea of the fundamental niche, determined by physiological properties of a species, more than 50 years ago. The idea remains both central to ecological thinking and largely unexplored experimentally. In this note, we describe some problems of using physiological experiments databases to estimate fundamental niches, show some solutions, and the apply our solution to testing the prediction that fundamental niches contain realized niches, for some species of marine fishes and terrestrial beetles. Our results were concordant with Hutchinson’s predictions.

## Introduction

In its modern form, the discipline called species distribution modeling (SDM) is based on the estimation of limits of tolerance to environmental variables, or ecological niche modeling (ENM) (Guisan & Zimmermann, 2000; Peterson et al., 2011). There is a relationship between geographical regions and their environmental features, called “Hutchinson’s Duality” (Colwell & Rangel, 2009) and this relationship enables estimating observed limits of tolerance in multivariate space, and then projecting them in geographical space, to obtain estimates of distributions.

In practice all the above can be done by software ignoring most of the conceptual framework. For instance, the package Maxent (Phillips, Anderson, & Schapire, 2006) simply asks for: (i) sets of coordinates of observations and, (ii) for sets of raster files representing environmental variables, and it performs the calculations that characterize the occurrence points in the multivariate environmental space. This is then are projected to geographic space. The entire procedure can be performed without mentioning the term “niche” even once.

However, several niche concepts are central to the interpretation of the procedures and results of most software used to estimate distributions. The basic idea is that of fundamental niche. A fundamental niche is the set of environmental combinations that produce a population fitness capable of sustaining viable populations (Drake, 2015; Holt, 2009; Hutchinson, 1957; Peterson et al., 2011) in the absence of biotic interactions. This is essentially a physiological concept, abstract and theoretical. In symbols, if *f*(***v***) is the function mapping environmental combinations into fitness, then the fundamental niche is **F**={**v** | *f*(***v***) > *0*}. The fundamental niche was originally conceived by Hutchinson as a simple rectangle in the multivariate environmental space, but soon people started using other shapes, for instance, ellipsoids (Brown, 1984; Drake, 2015; Maguire Jr, 1973), which somehow imply a structure (the expression in fitness units of regions of tolerance and their covariances) of fitness in environmental space. The fundamental niche is what evolves. In other words, the limits of tolerance and the covariations among them is what can be inherited and selected by evolutionary processes. The fundamental niche then deserves its name.

Now, notice that different regions of the world, and at different times, may have different environmental combinations. Therefore, we need a second niche concept, an “existing” fundamental niche, an idea introduced very briefly by Hutchinson(1957) and then elaborated by Jackson and Overpeck (2000). The existing niche is the fundamental niche actually existing in a given region, and at a given time. If the environments in a region **M**, at a time *t* are symbolized by **E**(**M**, *t*) = {**v** | **v** exists in region **M** at time *t*} then the existing niche is **F**\*(**M** *t*) = **E**(**M**, *t*)∩**F**. The existing niche is the combination of the physiological limits of tolerance expressed in a given region and time. It combines physiology with environment.

Finally, the environments in which a species can be observed are further limited by biotic interactions and by accessibility to movements. This defines the actual subset of the existing niche where a species can be observed. Any actual field observation will have to come from the existing niche restricted by availability, interactions and movements. The resulting set of actually observed environments is called the realized niche, denoted by **R**(**M**, *t*).

Hutchinson (1957) proposed a simple inequality relating the fundamental and the realized. Using the existing niche, and the symbols we defined above, the inequality is:

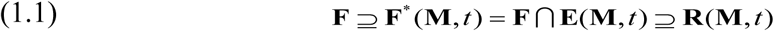

SoberÓn & Arroyo-Peña (2017) tested this inequality, but some of the subtleties of doing it were not well discussed. An extremely important question, the one we will explore here, is how can one combine physiological experimental data with climatic data, to establish the comparison in equation (1.1). This is by no means a simple operation (Addo-Bediako, Chown, & Gaston, 2000; Angilletta Jr & Angilletta, 2009; Bozinovic, Calosi, & Spicer, 2011; Kearney & Porter, 2009), mainly because physiological experiments are performed on simplified conditions, measuring variables that are not necessarily the ones that characterize environments in the field. To test equation (1.1) we will get data for the fundamental niches from a database of estimations of ranges of tolerance to extreme temperatures (Bennet et al. 2018), and data for the realized niches from occurrence reports, obtained from the Global Biodiversity Information Facility (GBIF; (Robertson et al., 2014).

The data required to estimate fundamental niches would be ideally derived from estimations of fitness under factorial experimental design. This is a task very seldom attempted, with the usual handful of exceptions (Birch, 1953; Hooper et al., 2008). Following Soberon & Arroyo-Peña (2017), we will use a single-variable (temperature) approximation to estimate the fundamental niche. We know that this is a very coarse approximation, but there is simply no data available on multivariate niche measurements using experiments. Even our single variable approach has some problems that we will discuss below.

## Methods

The objective of the present work is to illustrate some of the problems of attempting to relate temperature-tolerance data, obtained physiologically. To estimate fundamental niches we obtained data on ranges of temperature tolerance from the database GlobTherm https://datadryad.org/resource/doi:10.5061/dryad.1cv08/7 (Bennet et al. 2018). The ways they report extremes vary considerably between taxonomic groups, and some ways were unsuitable to estimation of fundamental niches. As examples, we included data from mammals, beetles and fish. For mammals, the Thermal Neutral Zone is reported, which is a range of temperature within which metabolic activity remains constant. This range is not really a good measure of the fundamental niche. For the beetles and fishes, 100% mortality lower and higher temperatures are reported. These are certainly compatible definition of a fundamental niche.

We also obtained information about the realized niche from records of species’ geographic occurrences. These were obtained from the Global Biodiversity Information Facility (GBIF; http://www.gbif.org). Validation of the taxonomic name of each species was done through consultation of The International Union for Conservation of Nature’s Red List of Threatened Species (IUCN; https://www.iucnredlist.org/) as well as other available biodiversity databases. We downloaded 1,739 points for mammals, 14,141 for fishes, and 3,558 for beetles. We subjected the data to a cleaning process by removing data fields that were either not relevant to our study or were incomplete or inconsistent (Costello & Wieczorek, 2014). This was done using Microsoft Excel and the R package ‘dismo’. We filtered data for cases with no georeference, occurrence records, repeated observations, records with no decimal precision, switches between longitude and latitude, changes in sign of geographic coordinates and using the program QGIS we were able to filter records outside the region of interest by mapping the occurrences in a world map raster.

Once this was done, we extracted bioclimatic variables for each species coordinates from WorldClim (https://www.worldclim.org). We used the annual mean temperature (bio1), maximum temperature of warmest month (bio5) and annual precipitation (bio12) in the spatial resolution of 10 minutes (∼340 km^2) which we then used to create one single raster file and then extracted the environmental values for each occurrence point using the R package ‘raster.’

To create an estimate of the accessible region we used the World Wildlife Fund map of the ecoregions of each continent (Olson et al., 2001) https://www.arcgis.com/home/item.html?id=67e8c7ce18f744f0b0e067c1e2247b6c). We extracted the ecoregion which matched the coordinates previously obtained for each species. We then assigned names and a code for each ecoregion in order to facilitate the manipulation of the data.

We then obtained a particular number of coordinates for both fish and beetles at random in order to reduce the sampling bias. In the case of the beetles species, there was a large difference between the number of samples found in the UK regions and the rest of the European continent (Spain, Belgium and the Netherlands). In order to remedy this we took 1000 samples at random from England and 1000 samples at random from the rest of the continent, thus taking into account the sampling bias that would otherwise interfere with the results. The same was done with the fish species.

We then used the gathered information with RStudio to fit a smooth histogram (a kernel density) to make a model of realized niche as well as the temperature in the M area for each species of fish and beetles. Hutchinson’s hypothesis is that the range of the fundamental would contain the observations (realized niche).

We also estimated the proportion of available environmental space that a particular species uses, by estimating the integral of the minimum between the black and green curves (Soberon & Arroyo-Peña, 2017).

## Results

The data on ranges of temperature tolerance obtained from the GlobTherm database were very contrasting in concept and in methodology. We found that they were not adequate to describe the fundamental niche in case of mammals, since GlobTherm reports a Thermal Neutral Zone (Bennet et al. 2018) which represents a range where metabolic activity does not change, therefore it encompasses an narrow and extremely favorable range, as we show in Figure 1, a random example out of the 61 cases of mammals in the database. All examples of mammals have very narrow TNZ ranges, often completely outside the actual observations.

**Figure 1.**
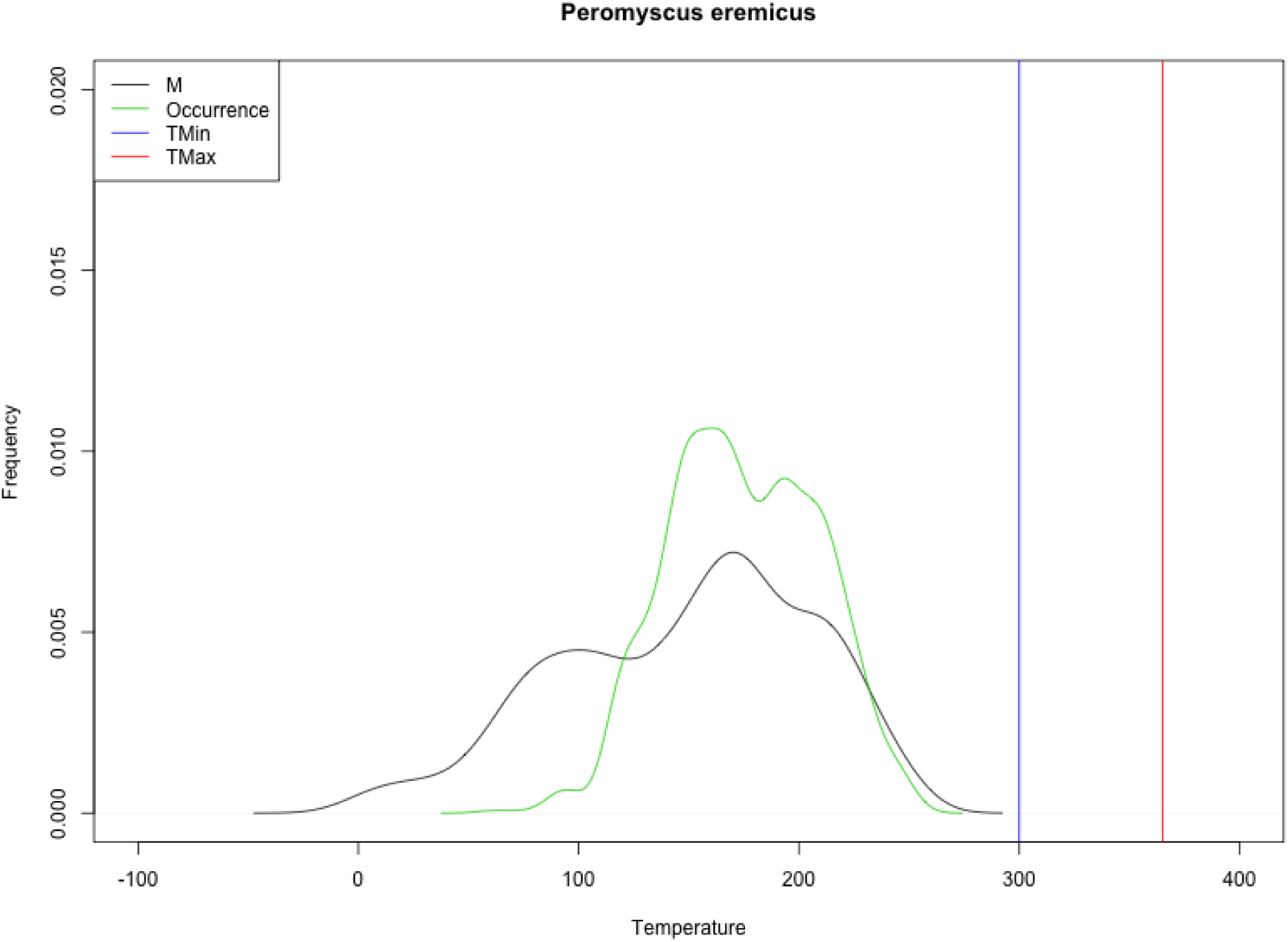
Range of Thermal Neutral Zone for one species of a desert mouse (blue and red lines), the smooth histogram of the available mean yearly temperature (times 10: BIO1 bioclimatic variable) in the M zone for *Peromyscus eremicus* (black line), and the smooth histogram of the GBIF observations (green line).

On the contrary, the data on ranges of temperature tolerance obtained from GlobTherm proved sufficient in the case of beetles and fish species, because these ranges are defined in terms of 50% and 100% mortality. These are able to portray a reasonable model for the fundamental niche for these species. In Figure 2 we present the data points used for the marine species, and in figure 3 the data records for the beetle species.

**Fig. 2.**
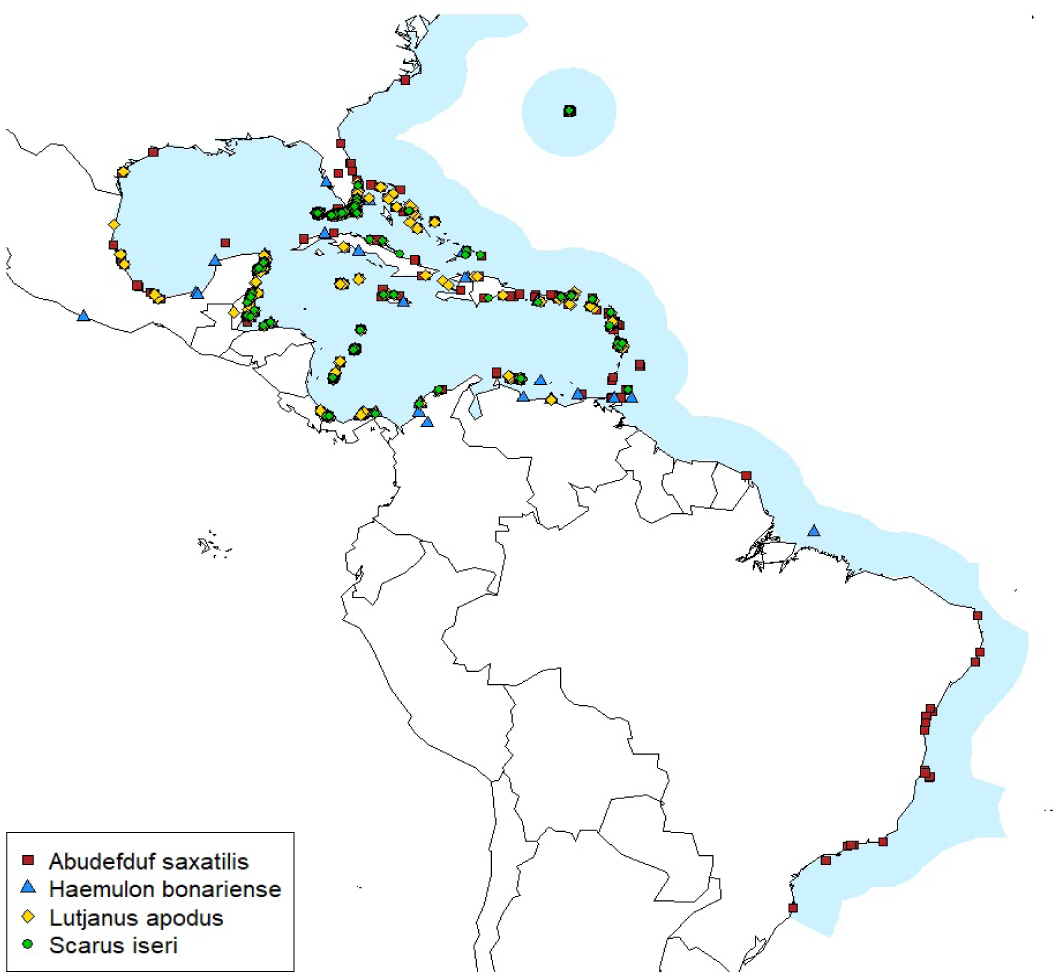
Map of occurrences of the fish species *<u>Abudefduf saxatilis</u>, <u>Haemulon bonariense</u>, <u>Lutjanus apodus</u>* and *<u>Scarus iseri</u>* along with the environmental availability region (M).

**Fig. 3.**
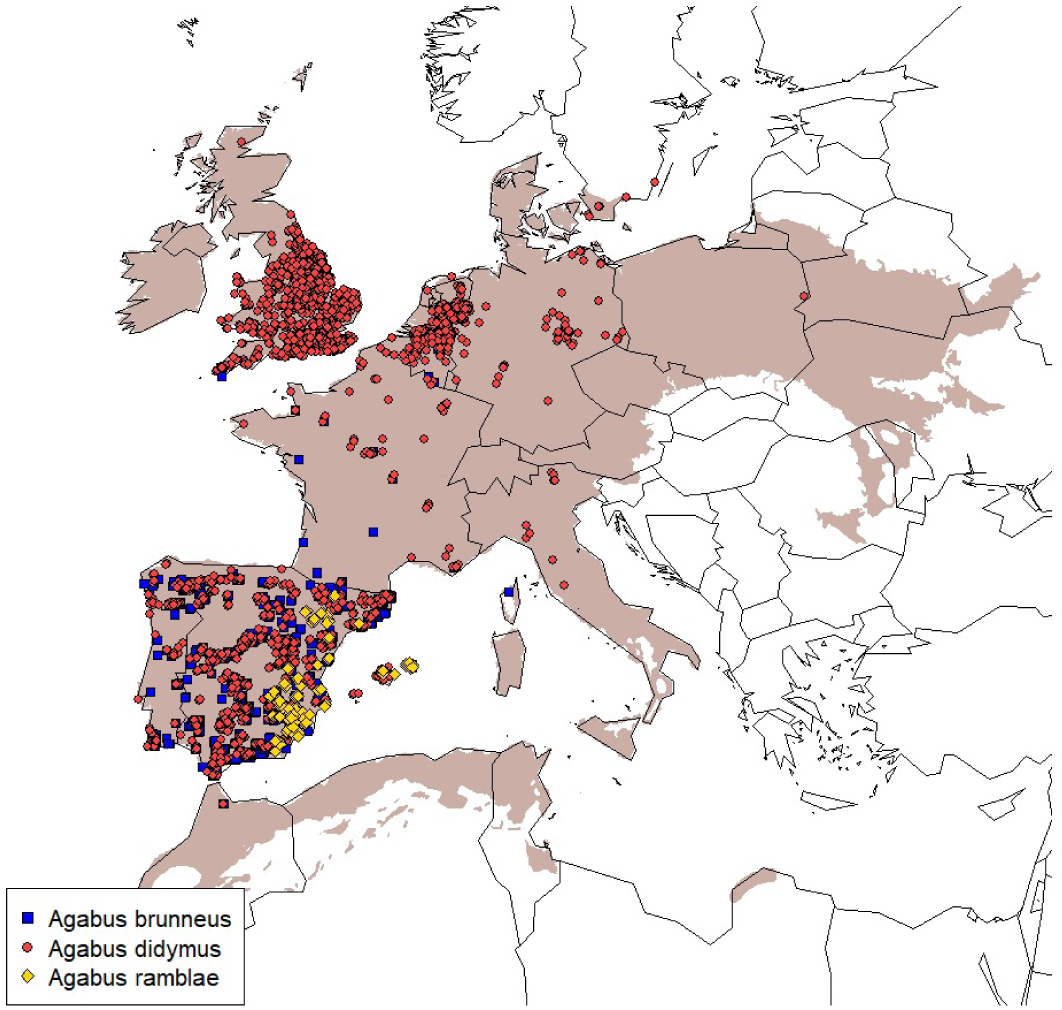
Map of occurrences of the beetle species *<u>Agabus brunneus</u>, <u>A. didymus</u>* and *<u>A. ramblae</u>* along with their environmental availability (M).

In figure 4-5 we present the results of niche measures for the fish and beetle species.

**Figure 4.**
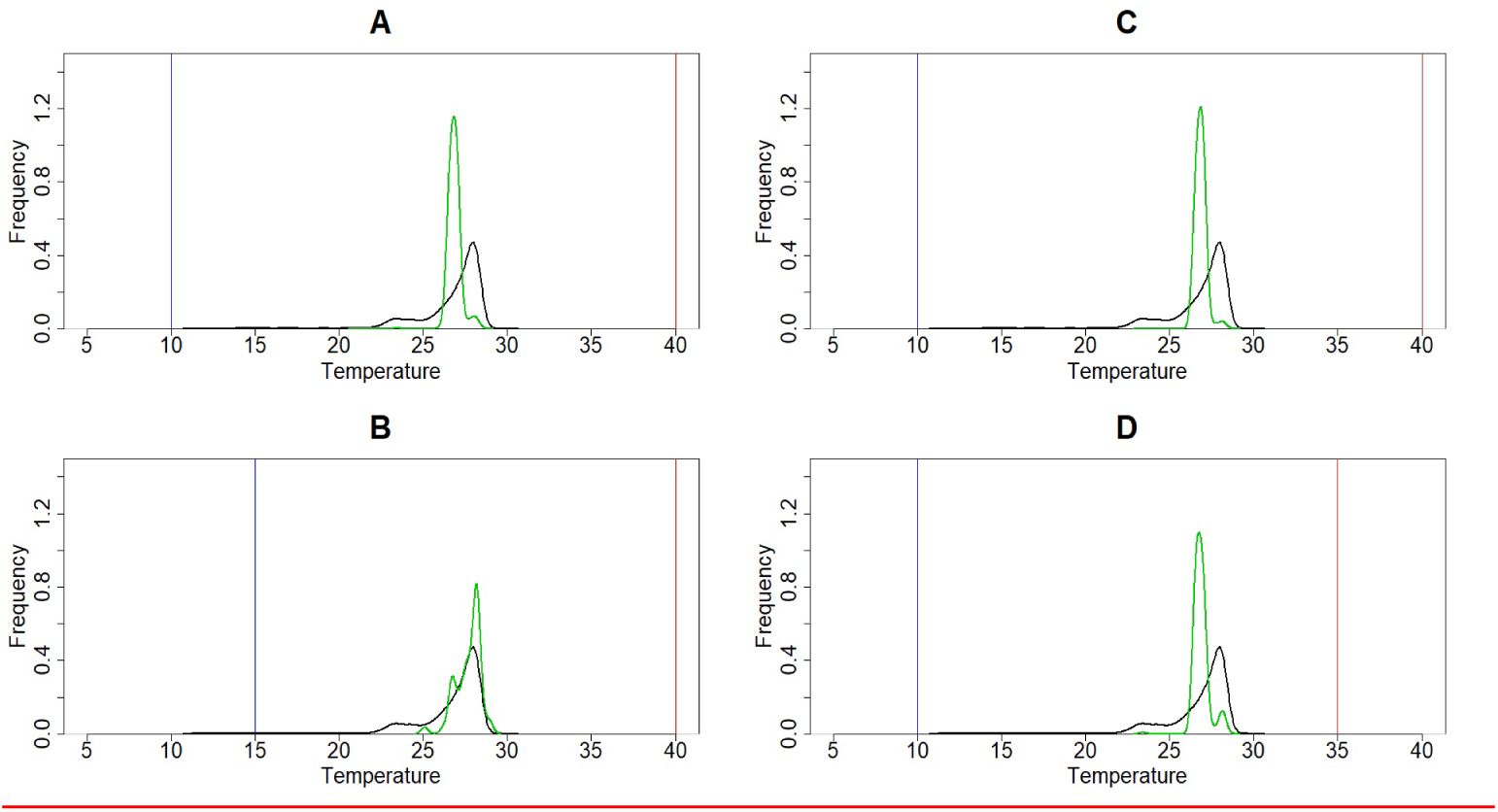
Graphs representing the fundamental niche, existent niche and realized niche of the species of fish *<u>Abudefduf saxati</u>* (A), *<u>Haemulon bonar</u>* (B), *<u>Lutjanus apodus</u>* (C) and *<u>Scarus iseri</u>* (D); *w*here the maximum temperature (red) and minimum temperature (blue) constitute the fundamental niche, the environmental availability M is the black line, and the number of species occurrences (green) is the realized niche.

**Figure 5.**
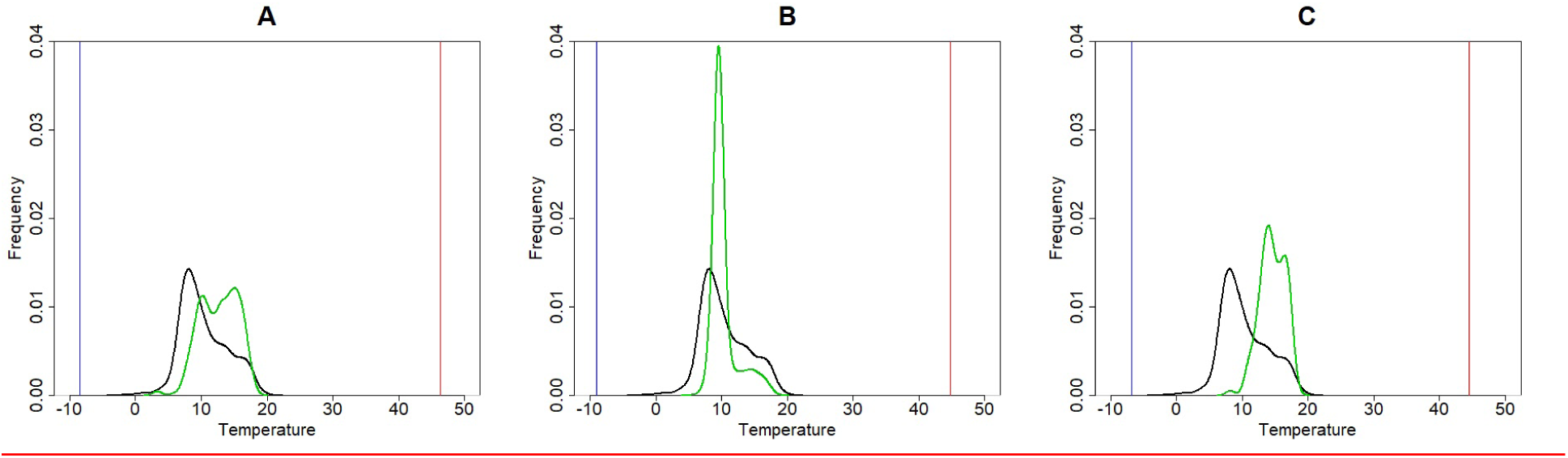
Graphs representing the fundamental niche, existent niche and realized niche of the species of beetle *<u>Agabus brunneus</u>* (A), *<u>A. didymus</u>* (B) and *<u>A. ramblae</u>* (C); where the maximum temperature (red) and minimum temperature (blue) constitute the fundamental niche, the environmental availability M (black) is the environmental availability in M and the number of species occurrences (green) is the realized niche.

Finally, in table 1 we present the proportion of environmental space occupied by the species, as estimated from occurrences from the GBIF database.

**Table 1.** Data and occupied niche for the species of beetles and fishes selected.

| Species | Tmax | tmin | Max Metric | Min Metric | Niche width | Occurrences | Clean Occurrences | Phylum | Class | Order | Family |
| --- | --- | --- | --- | --- | --- | --- | --- | --- | --- | --- | --- |
| *Agabus brunneus* | 46.4 | -8.5 | LT100 | LT100 | 0.92 | 1012 | 514 | Arthropoda | Insecta | Coleoptera | Dytiscidae |
| *Agabus didymus* | 44.8 | -9 | LT100 | LT100 | 0.51 | 5313 | 2885 | Arthropoda | Insecta | Coleoptera | Dytiscidae |
| *Agabus ramblae* | 44.5 | -6.8 | LT100 | LT100 | 0.95 | 114 | 114 | Arthropoda | Insecta | Coleoptera | Dytiscidae |
| *Abudefduf saxatilis* | 40 | 10 | LT100 | LT100 | 0.42 | 9989 | 5242 | Chordata | Actinopteri | Perciformes | Pomacentridae |
| *Haemulon bonariense* | 40 | 15 | LT100 | LT100 | 0.56 | 120 | 61 | Chordata | Actinopteri | Perciformes | Haemulidae |
| *Lutjanus apodus* | 40 | 10 | LT100 | LT100 | 0.27 | 9987 | 6872 | Chordata | Actinopteri | Perciformes | Lutjanidae |
| *Scarus iseri* | 35 | 10 | LT100 | LT100 | 0.44 | 9997 | 1796 | Chordata | Actinopteri | Perciformes | Labridae |

## Discussion

An important lesson from our results is that data of tolerance curves, even for the simplified case of a single variable, cannot be simply assumed to represent a fundamental niche in the sense of Hutchinson. Hutchinson (1957) defined the niche in terms of fitness, and unless the experiments have a direct relationship to fitness, their use as proxies for fundamental niches is problematic. The Thermal Neutral Zone for the endotherms, as reported in the GlobTherm database (Bennet et al. 2018) fail to fulfill this criteria, and should not be used as a proxy for the range of positive fitness. On the other hand, the 100% mortality critical temperatures are indeed represent a fitness range, albeit a bit extreme.

Using the ranges compatible with the concept of fundamental niche, Hutchinson’s inequality is proved, adding confirmatory evidence to previous results by Soberon & Arroyo-Peña (2017). In all the examples we present estimating the fundamental niche with the 100% critical range of temperatures, the entire available range of environments occurs inside the fundamental niche. This means that the entire region of hypothetical availability (M) could be occupied. This is certainly not the case, and there are several reasons why there may be empty regions of accessible and favorable space. The first one was proposed by Hutchinson (1957): maybe there are negative interactors (competitors or predators) preventing the expansion of the range of a species. The second one may be the result of using a single variable as a proxy to a multivariate niche. Other variables may be limiting the actual distribution of the species, but we are measuring just one. By adding variables one add constraints to a niche function (Soberón & Arroyo-Peña, 2017), and therefore reduces the proportion of available environmental space used. The third reason is very similar to the second: unoccupied favorable temperatures may be due to lack of suitable habitat. Habitat, as opposed to climate, may be a limiting factor. For instance, in the case of the freshwater beetles *Agabus*, climate may be favorable but if the correct type of freshwater body is lacking, the species may not be present. The value of the integrals in table 1 represent the proportion of available climate actually used by the species (assuming the GBIF data is representative).

The above point illustrates the need to have a hypothesis about accessible geographic space, or M (Barve et al. 2011). An M hypothesis is not only indispensable when using software sensitive to the background data, like Maxent (VanDerWal, Shoo, Johnson, & Williams, 2009), but also to interpret the results of any niche modeling. Knowledge about the amount of available niche space suggests whether missing factors are present.

The work we present here should be regarded as an exploration into the problems of estimating fundamental niches. The main problem is the lack of experimental data, relating many relevant variables to fitness values. This is an obvious problem, and it is very puzzling to notice how little, comparatively, experimental work has been done to address it. Besides temperature physiological ranges, only a handful of experiments include two variables (Birch, 1953; Hooper et al, 2008), and none, to our knowledge, more than 2. This represents an open field of research for the future.

